# Italian universities do not recruit according to proficiency in scientific production

**DOI:** 10.1101/2021.10.25.465694

**Authors:** Pasquale Gallina, Berardino Porfirio

## Abstract

We analyzed the files regarding recruitment competitions won by 186 professors of selected bibliometric disciplines in Florence between January 2014 and 30 June 2021. An equal number of professors recruited at other Italian universities and of researchers who never attained professorship in Italy were randomly chosen in the same disciplines as each Florentine professor among individuals possessing National Scientific Qualification, a prerequisite for professorship. The H-indexes at the time of qualification (T1), of the Florence call (T2), and the current (T3) time were obtained from Scopus.

Non-recruited researchers were more likely (Chi-square test) to show a higher H-index than both Florentine (T1 p=0.0005, T2 p=0.0015, T3 p=0.0095) and non-Florentine professors (T1 p=0.0078, T2 p=0.0245, T3 p=0.0500). Fifty-four non-recruited scientists serve in foreign universities, 100 at national/international research centers. The remaining 32 scientists (25 who keep producing despite precarious employment, and seven who have stopped publishing) were, at any rate, as likely as Florentine (T3 p=0.69) and non-Florentine professors (T3 p=0.14) to show a higher H-index.

This study suggests that Italian academia does not recruit professors according to their qualitative/quantitative ability to publish, a detriment to knowledge for the nationwide system and on a global scale.

## Introduction

Malpractice in the system of recruitment at Italian universities has been reported (Assad, 2016; Rigante, 2016). Academia is plagued by nepotism (Grilli and Allesina, 2017), clientelism and deference to politics (Gallina and Gallo, 2020). Calls are in most cases tailored to favor local candidates, with a consequential lack of mobility and independence (Gallina and Gallo, 2020). Many worthy scientists, who are not close to influential figures, remain marginalized and have either had to leave Italy to continue their career or give up on their ambitions. An impoverishment in scientific production for Italy is expected to follow. Here, we aim to demonstrate this issue in a quantitative manner.

## Methods

From the archives of the University of Florence (https://www.unifi.it/vp-2456-docenti-e-ricercatori-di-ruolo.html), we retrieved the files regarding recruitment of 186 professors in selected bibliometric disciplines, according to law 240/2010 (Presidente della Repubblica, 2011) since its inception until June 30, 2021.

### Search strategy

With reference to Ministerial Decree 29 July 2011 n. 3361 (Ministro della Pubblica Istruzione, 2011) until 2015, and to Ministerial Decree 30 October 2015 n. 8552 (Ministro della Pubblica Istruzione, 2015) thereafter, we identified the competition sectors composed of a single academic discipline in the fields of Social Sciences and Humanities, Physical Sciences and Engineering, and Life Sciences according to the European Research Council (ERC - https://erc.europa.eu). This initial step was undertaken to identify in the subsequent matching phase (see below) researchers belonging, with certainty, to the same academic disciplines.^1^ Following a previously described method (Gallina and Gallo, 2020), in the time window April-June 2021 we identified the recruitment procedures in the above selected academic disciplines, according to art. 18, commas 1, 4 and art. 24, commas 5, 6 of law 240/2010 (Presidente della Repubblica, 2011), held in Florence between January 2014 and 30 June 2021; for each competition, the winner was identified. We then looked for the year (the quarter for the 2016-2018 session) when each of the recruited professors in Florence achieved National Scientific Qualification, a pre-requisite for professorship as per art. 16, law 240/2010 (Presidente della Repubblica, 2011), by searching their competition sectors in the *Abilitazione Scientifica Nazionale* website (https://abilitazione.miur.it/public/index.php). Among individuals who achieved the National Scientific Qualification along with each of the recruited Florentine professors, we randomly identified two scientists: one employed at other Italian universities and another who never achieved an academic position in Italy (https://cercauniversita.cineca.it/php5/docenti/cerca.php). For the 2016-2018 session of National Scientific Qualification, when the quarter in which the Florentine professor achieved qualification did not contain non-recruited researchers, the search was extended to the nearest quarters. When drawing resulted in duplicates, the procedure was repeated. In three instances, a suitable non-recruited researcher was not found, thus the entire data string was omitted. Information about the current work place of non-recruited researchers was obtained from the affiliation stated in their last published article as reported on Scopus (https://www.scopus.com/search/form.uri?display=authorLookup#author) and reasonably confirmed by an internet search. The most recent year of publication for the non-recruited researchers was obtained from the same source. For all scientists, H-indexes at the time of their National Scientific Qualification (T1) and at the time of the Florence call (T2) were retrieved from Scopus (https://www.scopus.com/search/form.uri?display=authorLookup#author). The current H-index (T3) was obtained from the same source in the time-window 1 - 31 July 2021. Before combining records, disambiguation among authors was obtained through verification of researcher IDs and/or CVs.

### Statistics

We posited that, among the population of those who achieved National Scientific Qualification, the probability that a random draw within a certain competition sector (i.e. within a single academic discipline) would result in a person with an H-index higher (or lower) than that of the Florentine winner is 50%. We determined this probability assuming that the National Scientific Qualification is fair, i.e. “heads or tails” are equally likely once ties are removed. The null hypothesis, i.e. that for a researcher who obtained National Scientific Qualification having an H-index higher (or lower) than that of another person drawn from the same pool is equal, was tested using the Chi-squared distribution with 1 degree of freedom according to the formula (H-L)^2^/(H+L), where H and L are the number of higher and, respectively, lower outcomes. P-values <0.05 were considered evidence of biased deviation from the expected H=L.

### Sensitivity analyses

Contrary to the criteria used in the primary analysis where individuals employed at other Italian universities as assistant, associate or full professors were matched with the Florentine professors, we performed a second series of drawings to check the robustness of the results in which only associate and full professors were considered. This assumes that those who were appointed as associate or full professors would perform better (on average) than assistant professors, thus providing a more stringent scenario when comparing their H-indexes with those of both the Florentine professors and the non-recruited individuals.

We decided to use Scopus because it offers a more extensive list of modern sources, which seemed best suited to our investigation. However, we retrieved the H-indexes for each of the 186 triplets at each of the three measurement times also from Web of Science (https://www.webofscience.com/wos/author/search) to ascertain whether different sources and the way information is collected in highly heterogeneous fields may affect results.

## Results

Chi-square test showed that the probability for a Florentine professor to have an H-index higher than that of a non-Florentine counterpart was not different from 50% (T1: 87 *vs* 86, 13 ties, p=0.94; T2: 78 *vs* 93, 15 ties, p=0.25; T3: 81 *vs* 90, 15 ties, p=0.49). On the other hand, non-recruited researchers were more likely to show a higher H-index than both Florentine (T1: 109 *vs* 63, 14 ties, p=0.0005; T2: 108 *vs* 66, 12 ties, p=0.0015; T3: 103 *vs* 69, 14 ties, p=0.0095) and non-Florentine professors (T1: 104 *vs* 69, 13 ties, p=0.0078; T2: 104 *vs* 74, 8 ties, p=0.0245; T3: 101 *vs* 75, 10 ties, p=0.0500). Most non-recruited researchers serve in foreign universities (n=54) or national/international research centers (n=100). Some others (n=25) continue producing despite precarious employment, whereas seven scientists have stopped publishing, as suggested by at least three years of inactivity. Notwithstanding, this subset of 32 non-recruited researchers were, at any rate, as likely as Florentine (T3: 12 *vs* 14, 6 ties, p=0.69) and non-Florentine professors (T3: 11 *vs* 19, 2 ties, p=0.14) to show a higher H-index.

Sensitivity analyses confirmed the above results.

## Discussion

The aim of the present study was to demonstrate if, within a same discipline, professors recruited in Italy performed better or worse in terms of scientific production, as compared to scientists who have not accessed Italian academia. The H-index represents the most straightforward metric tool because of its combination of productivity and impact indicators, although the value of bibliometric data as exhaustively indicating the merits of a researcher is debated (Gasparyan et al., 2018; Braithwaite et al., 2019) and it is undisputed that teaching and managerial abilities are also required to hold academic positions.

With these premises, this study suggests that the Italian academia disregards meritocracy, at least in terms of qualitative and quantitative ability to publish. Paradoxically, if the Florentine recruitment had been approached randomly, rather than according to bylaw procedures, a corpus of professors with higher H-indexes on average would have been selected. In fact, about 30% of the individuals who achieved qualification in the first session held in 2012 were not on university payrolls in Italy (Gallo, 2014). Since non-recruited researchers are more likely to perform better in bibliometric terms, the nationwide system is expected to suffer a detriment (Abramo and D’Angelo, 2018), if only because they were educated with public resources. Studies should quantify the amount of knowledge generated by Italian education and lost by our academic system in terms of GDP reduction. Fortunately, this human capital is not completely lost. Almost 83% of these individuals are serving foreign universities or national/international research centers. However, in giving up academic aspiration, a not negligible proportion of valued, rejected researchers deprive science of their potential contribution on a global scale.

It is essential for the Italian ministerial authorities to investigate the aberrant mechanism that leads to the exclusion of qualified researchers from universities. However, for there to be real change the crucial role of three actors cannot be ignored: the Italian academic establishment must carry out a profound self-criticism of its work so far; the world scientific community, impoverished by this malpractice, must operate a moral suasion on Italian universities and support the efforts of people who fight against this cancer in our country; early-career Italian researchers must embrace the call to create a more equitable, collaborative and healthy academia (Aguilar, 2021) and avoid complicity.

### Notes

1. Competition macro areas (*macro settori*) correspond to ERC search domains. These are subdivided in competition sectors (*settori concorsuali*), which correspond to ERC panels. The latter can involve a single academic discipline (*settore scientifico disciplinare*), corresponding to ERC panel descriptors, or be split into more than one of these latter.

## Research data

We are willing to make available all anonymized data upon request to the corresponding author.

